# Ethanol Exposure Perturbs Sea Urchin Development and Disrupts Developmental Timing

**DOI:** 10.1101/2022.07.07.499183

**Authors:** Nahomie Rodríguez-Sastre, Nicholas Shapiro, Dakota Y. Hawkins, Alexandra T. Lion, Monique Peyreau, Andrea E. Correa, Kristin Dionne, Cynthia A. Bradham

**Affiliations:** Biology Department, Boston University, Boston MA; Bioinformatics Program, Boston University, Boston MA; MCBB Program, Boston University, Boston MA; Universidad de Puerto Rico-Recinto Aguadilla, Puerto Rico; Biological Design Center, Boston University, Boston MA

**Keywords:** ethanol, sea urchins, Hedgehog, Retinoic Acid, Timing, FAS

## Abstract

Ethanol is a known vertebrate teratogen that causes craniofacial defects as a component of fetal alcohol syndrome (FAS). Our results show that sea urchin embryos treated with ethanol similarly show broad skeletal patterning defects, potentially analogous to the defects associated with FAS. The sea urchin larval skeleton is a simple patterning system that involves only two cell types: the primary mesenchymal cells (PMCs) that secrete the calcium carbonate skeleton and the ectodermal cells that provide migratory, positional, and differentiation cues for the PMCs. Perturbations in RA biosynthesis and Hh signaling pathways are thought to be causal for the FAS phenotype in vertebrates. Surprisingly, our results indicate that these pathways are not functionally relevant for the teratogenic effects of ethanol in developing sea urchins. We found that developmental morphology as well as the expression of ectodermal and PMC genes was delayed by ethanol exposure. Temporal transcriptome analysis revealed significant impacts of ethanol on signaling and metabolic gene expression, and a disruption in the timing of GRN gene expression that includes both delayed and precocious gene expression throughout the specification network. We conclude that the skeletal patterning perturbations in ethanol-treated embryos likely arise from a loss of temporal synchrony within and between the instructive and responsive tissues.

## Introduction

Pattern formation during embryonic development represents the expansion of genetically encoded biological design into tangible physical structures within the animal (Briscoe, 2019; Cang et al., 2021; Ulloa and Briscoe, 2007; Zernicka-Goetz, 2002). Studying patterning in vertebrates is challenging, because of both their morphological and their genomic complexity. Vertebrates exhibit two ancestral genome duplication events that have resulted in significant genetic redundancy. Sea urchin embryos offer a considerably simpler model for pattern formation. First, they are morphologically quite simple, and second, because their genome was never duplicated, they lack extensive genetic redundancy (Sodergren et al., 2006), simplifying the study of gene function in this model.

The sea urchin larval skeleton is a bilaterally symmetric biomineral composed of calcium carbonate with numerous embedded proteins; the skeletal elements are secreted by the primary mesenchyme cells (PMCs) (Mann et al., 2010; Wilt, 2002) The PMCs ingress into the blastocoel at the onset of gastrulation, then migrate into stereotypical positions (Lyons et al., 2012). The PMCs receive instructive cues, mainly from the ectoderm, that direct their spatial positioning and differentiation (Adomako-Ankomah and Ettensohn, 2013; Armstrong et al., 1993; Duloquin et al., 2007; McIntyre et al., 2013; Piacentino et al., 2016a, 2016b, 2015; Walton et al., 2009). The PMCs migrate into a primary (1°) spatial arrangement at late gastrula stage that is comprised of a posterior ring of cells around the blastopore with ventrolateral PMC clusters that extend cords of PMCs towards the embryo’s anterior pole. This ring-and-cords arrangement presages the 1° skeletal pattern (Fig. 1A, blue). Then, additional PMC migration produces the secondary (2°) elements that give rise to the pluteus skeleton (Fig. 1A, red). Previous work from our lab and others has discovered numerous skeletal patterning cues that are expressed by the ectoderm; the molecules that drive skeletal patterning include conserved signaling proteins, second messengers, and extracellular matrix molecules (Armstrong et al., 1993; McIntyre et al., 2013; Piacentino et al., 2016b, 2016a, 2015). Although most patterning cues are expressed by the ectoderm, Hedgehog (Hh) signals from the endoderm also contribute to skeletal patterning (Walton et al., 2009).

**Figure 1.**
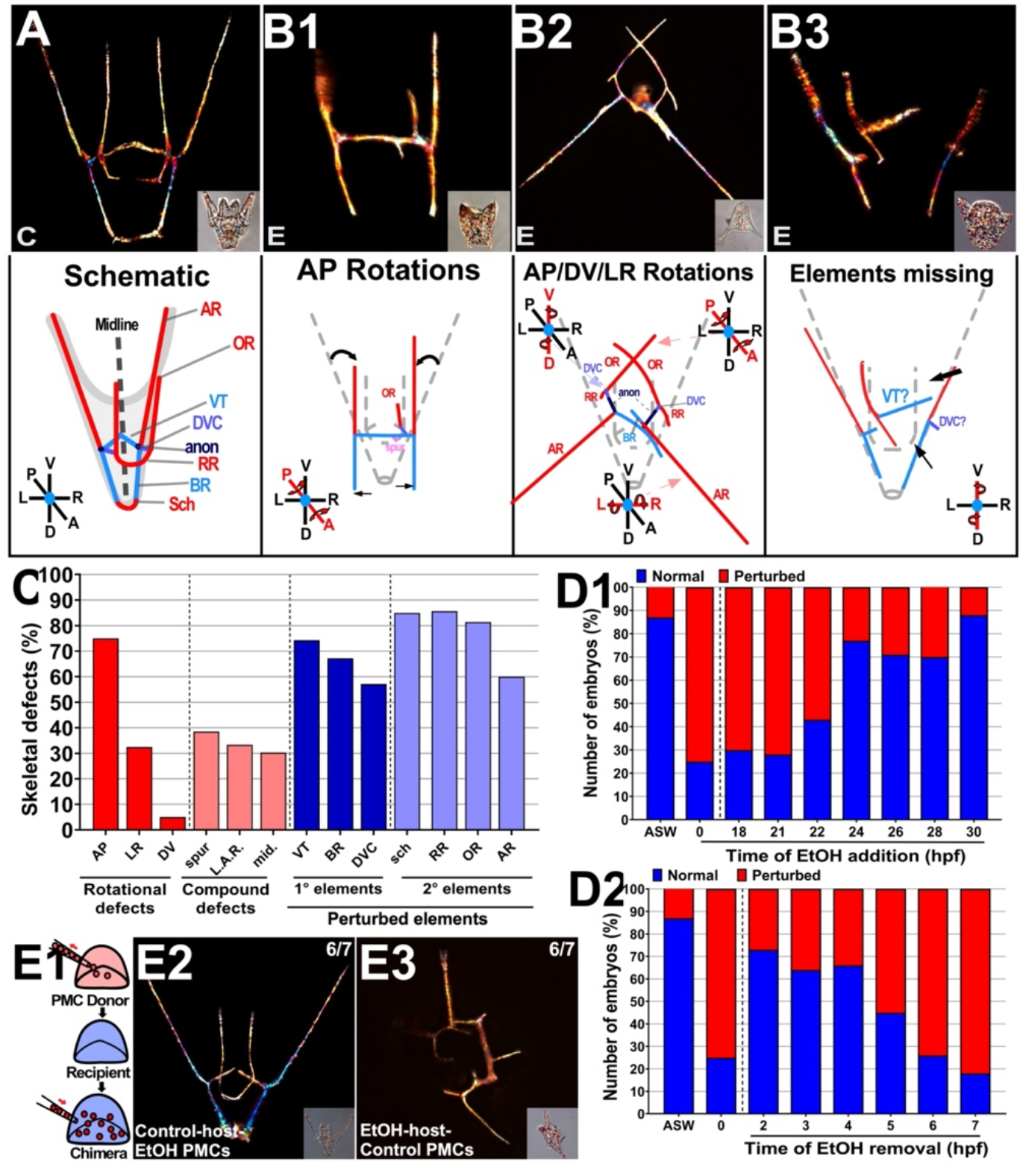
Ethanol exposure perturbs skeletal patterning between 5 and 22 hpf. A-B. Representative skeletal morphology (birefringence, top) and schematics (bottom) of control (C, A) and EtOH-treated (E, B1-3) embryos are shown at 48 hpf (pluteus stage); the corresponding DIC images are inset, and body axes are indicated. The primary skeletal elements (A, blue) include the initial triradiate that gives rise to the ventral transverse rods (VTs), the dorsoventral connecting rods (DVCs), and the anonymous rods (anon); the latter branch to produce the dorsal body rods (BRs). Secondary elements (A, red) include the posterior aboral rods (ARs) that branch from the anons, as well as the anterior oral rods (ORs) and the recurrent rods (RRs) that branch from the DVC, and finally, the scheitel (sch) that branches from the tips of the BRs at the dorsal extreme. The perturbed skeletons in EtOH-treated embryos exhibit abnormal rotations of skeletal elements about one of the body axes (B1), sometimes in combinations (B2), as well as element losses (B3). Arrows show rotations (B1, B2) and absent elements (B3). C. Skeletal patterning defects, including rotational defects, compound defects, and element losses, are plotted as the percentage of embryos exhibiting the indicated effect. Spur, spurious elements; LAR, long anonymous rods; mid, midline losses; n > 60. D. The percentage of embryos that were normal (blue) or perturbed (red) after adding (D1) or removing (D2) EtOH at different time points is shown; n > 50 embryos per time point. Artificial sea water (ASW) control and EtOH controls (treated at 0 without removal) are the same in each plot. E. The approach for PMC transplantation to produce chimeric embryos is shown schematically (E1). Chimeric embryos are shown as morphology (DIC, 1) and skeletal pattern (birefringence, 2) for EtOH-treated PMCs in control hulls (E2) or the reciprocal control PMCs in EtOH-treated hulls (E3) at 40 hpf (pluteus stage). The results shown were obtained in 6/7 trials. Embryos were treated with EtOH at fertilization (0 hpf) then removed from EtOH prior to microsurgery at mesenchyme blastula stage, following PMC ingression (at approximately 12 hpf). See also Fig. S1.

Ethanol (EtOH) exposure during embryonic development in vertebrates leads to Fetal Alcohol Spectrum Disorder (FASD), which is a group of conditions that include a range of neurological and skeletal patterning defects (CDC, 2021; Eberhart et al. 2016; Kot-Leibovich and Fainsod, 2009a). Fetal Alcohol Syndrome (FAS) in humans is characterized by defects in the central nervous system (CNS) and facial aberrations (O’Neil, 2012). The patterning defects that arise after EtOH exposure primarily affect the midline of the face and are thought to arise from perturbations to the neural crest cells that produce the facial skeleton (Abramyan, 2019; Cartwright and Smith, 1995; Smith et al., 2014). Studies performed in vertebrate models have identified multiple downstream targets for EtOH, including the Hh signaling pathway (Abramyan, 2019; Ahlgren et al., 2002; Li et al., 2007) and the retinoic acid signaling pathway (Hong and Krauss, 2017; Kot-Leibovich and Fainsod, 2009; Shabtai et al., 2018). Although these pathways contribute to the FASD phenotype, the overall mechanism underlying the FASD phenotypes, including their variable penetrance and severity, is still not well understood (Eberhart et al. 2016; Sarmah et al. 2020; Serio et al. 2019).

We performed a series of experiments to understand whether EtOH mediates FAS-like effects in sea urchin embryos, which offer a less complex model system and, as basal deuterostome, provide a compelling evolutionary comparison to vertebrates. When we exposed sea urchin zygotes and embryos to EtOH, we observed skeletal patterning defects, including midline and rotational defects, which could be considered analogous with facial patterning defects in vertebrate FAS. In contrast, EtOH-treated sea urchin zygotes did not exhibit neural losses, unlike EtOH-exposed vertebrate embryos. Surprisingly, we found that RA or Hh signaling perturbation does not account for the EtOH phenotype in sea urchins. We found that migration of the skeletogenic PMCs was delayed in EtOH-treated embryos, as was the expression of some PMC-specific or -enriched genes and ectodermal patterning cues. Temporal transcriptome analysis revealed that other such genes exhibit precocious expression, indicating that ethanol treatment results in temporally disrupted developmental gene expression; this broad asynchrony may underlie the defects in the intercellular communication-based skeletal patterning process in sea urchin embryos.

## Methods

### Reagents

Dose-response experiments were performed to determine the optimal working doses for each perturbation reagent, which include EtOH, Acetaldehyde, Retinol, Retinoic Acid, Fomepizole (Sigma-Aldrich or Fisher Scientific), Cyclopamine (Enzo Life Sciences), and SAG (Santa Cruz Biotechnology). All other chemicals were obtained from Sigma-Aldrich or Fisher Scientific unless otherwise noted.

### Chemical Treatments

Embryos were cultured in artificial sea water (ASW) at a density of 500 embryos/mL in a 24 well plates and incubated in humid chamber at 23°C. EtOH- and Acetaldehyde-treated embryos were cultured in a separate well-plate and humid chamber to prevent vapor-mediated cross-contamination. Highly volatile and reactive retinol was reconstituted in DMSO, aliquoted, and added to embryo cultures inside an oxygen-free glove box.

### Animals, perturbations, transplant, imaging, and skeletal scoring

*Lytechinus variegatus* sea urchins were obtained from the Duke University Marine Laboratory (Beaufort, NC) or from Reeftopia (Miami, FL). Gamete harvesting, fertilizations, and embryo culturing were performed as previously described (Bradham et al., 2006). Transplant experiments were performed as described (Piacentino et al., 2016b), using donor embryos labeled with tetramethyl rhodamine methyl ester (TMRM, Fisher Scientific). To produce contemporaneous stage-matched embryos for PMC transplants, control embryos were fertilized 30 minutes prior to embryos that were treated with EtOH to account for delayed PMC ingression with EtOH treatment. Fluorescent labeling of embryos was performed 30 minutes before the removal of PMCs. EtOH treatment was from fertilization to mesenchyme blastula stage (MB), at approximately 12 hpf. Embryos were washed from EtOH at MB, prior to performing PMC swaps. Larval morphology was imaged using DIC; larval skeletal birefringence was imaged in multiple focal planes using plane-polarized light on a Zeiss Axioplan inverted microscope at 200x magnification. Focal planes were then manually assembled into montage images using CanvasX (Canvas GFX, Inc.) to present the complete larval skeleton in focus. All focal planes were used for scoring with our in-house scoring rubric (Piacentino et al., 2016b) which captures element shortening, lengthening, loss, or duplication, spurious element production, abnormal element orientations, as well as whole embryo-level defects such as midline defects, and orientation defects.

### Immunostaining and confocal microscopy

Immunolabeling was performed as described (Bradham et al., 2006). Primary antibodies were PMC-specific 6a9 (1:5; from Charles Ettensohn, Carnegie Mellon University, Pittsburgh, PA), neural-specific 1e11 (1:10, from Robert Burke, University of Victoria, BC, Canada) and anti-serotonin (Sigma), and ciliary band-specific 295 (undiluted; from David McClay, Duke University, Durham, NC). Confocal imaging was performed using an Olympus FV10i laserscanning confocal microscope. Confocal z-stacks were projected using Fiji, and full z-projections are presented.

### Fluorescent *in situ* hybridization (FISH)

Full-length probes for LvJun, LvVEGF, LvVEGFR, LvHh, and LvWnt5a were transcribed using SP6 or T7 RNA polymerases (New England BioLabs) and labeled with digoxigenin (Roche). *In situ* hybridization was performed as previously described (Piacentino et al., 2015). FISH probe for Hedgehog, VEGF, VEGFR, and Wnt5 were previously described (McIntyre et al., 2013; Piacentino et al., 2015; Walton et al., 2009).

### HCR FISH: probe sets, amplifiers, and buffers

Embryos were collected and fixed in 4 % paraformaldehyde at 18 hours post-fertilization (hpf). The published hybridization chain-reaction (HCR) single molecule FISH protocol (Choi et al., 2018, 2016) was performed using fluorescently labeled amplifiers (hairpins with either Alexa488, Alexa 546, or Alexa647 labels), buffers, and probe sets from Molecular Instruments, Inc. (Los Angeles, CA). The detection step was performed in 0.5 mL microcentrifuge tubes at 37° C; embryos were then transferred to a 96-well plate for the amplification step. Embryos were incubated in the hairpin solution for ~2.5 hours or overnight in the dark at room temperature, washed with 5X SSCT, then mounted in PBS/glycerol for imaging. Probe sets were designed from the open reading frames by Molecular Instruments, Inc. (Los Angeles, CA).

### Acetaldehyde Measurements

Embryos were cultured at a density of 1500 embryos/mL until18 hpf, when the culture supernantant was collected. Acetaldehyde measurements was performed with a kit (Megazyme, Inc.) according to the manufacturer’s instructions.

### Serotonin level and spatial gene expression measurements

Fiji was used to process all confocal z-stacks. For measurements of serotonin levels from immunostained images, regions of interest (ROIs) were manually drawn in Fiji around each neural cell body, then the fluorescence intensity was measured as the total relative fluorescent units (RFU) per cell. Comparisons were limited to controls and treated embryos that were prepared together and imaged using identical settings. For spatial measurements of gene expression territories from FISH images, a threshold was applied to z-projection images in Fiji to produce binary images. ROIs were then defined in an automated manner using the Analyze Particles function in Fiji after adjusting the size range to most accurately capture the data. The resulting area values were then normalized to the total area of the z-projected embryo.

### RNA-seq and Analysis

Total RNA was prepared from 10,000 embryos per sample using TRIzol (Invitrogen) and precipitated along with glycogen carrier (Ambion) from control and ethanol-treated embryos at 15, 18 and 21 hpf, from three independent biological replicates. Library preparation and transcriptome sequencing were performed using DNA nanoball (DNB)-seq (BGI, Inc.) to generate 100 bp paired end reads. Quality control was performed using fastp to trim low-quality bases and remove low quality and low complexity reads (Chen et al., 2018). Reads were then aligned to the *Lvar 3.0* genome (Davidson et al., 2020) using *STAR* (Dobin et al., 2013) and read counts were calculated using *featureCounts* (Liao et al., 2014). Raw fastq files and the processed count matrix are available at GEO (accession number GSE207100). Multiple gene models mapping to a single gene annotation were collapsed to a single entry by summing read counts across models. Differential expression analysis and library normalization was performed using DESeq2 (Love et al., 2014), with batch set as a covariate during differential expression analysis. For downstream analysis, normalized counts were batch corrected using COMBAT from the *sva* R package (W. E. Johnson et al., 2007; Leek et al., 2012). Principal component analysis was performed using the *prcomp* R function with the corrected counts. GO enrichment analysis was performed using ssGSEA from the *gsva* R package (Barbie et al., 2009; Hänzelmann et al., 2013), and the top 20 up and down pathways were ranked by p-value following a Wilcoxon Rank-sum test. The top 20 enriched and depleted GO terms were manually binned using custom-defined categories (Hogan et al., 2020). Gene regulatory network (GRN) and skeletal patterning cue gene expression levels were analyzed in time-matched and heterochronic comparisons; in time-matched comparison, genes were considered to be early/elevated or late/reduced if the average expression level in EtOH-treated embryos deviated from control embryos by ≥ 18%. Potential GRN circuits connecting genes whose expression was similarly impacted by EtOH were manually mapped onto known and predicted edges in an integrated version of the specification network (Hogan et al., 2020).

## Results

### EtOH treatment results in skeletal patterning defects

To test whether EtOH has teratogenic effects on developing sea urchins, we exposed zygotes to a range of EtOH doses, then assessed their larval phenotypes. Using a systematic scoring approach (Piacentino et al., 2016b), we found that treatment with 1.7% (369 mM) EtOH induces dramatic skeletal patterning defects that include element losses, spurious elements, and rotational defects, with anterior and ventral elements most frequently perturbed, and both 1° and 2° elements affected (Fig. 1A-C). Most skeletal elements exhibited losses in EtOH-treated embryos (Fig. 1C). Anterior-posterior rotational defects were the most common orientation defect in EtOH-treated embryos. The treated embryos also showed unusual defects, including a high frequency of spurious elements, long anonymous rods, and midline losses (Fig. 1C); these latter defects were not observed with perturbations to other known patterning cues or their receptors, including Univin/Alk4/5/7, VEGF/VEGFR, SLC26a2/7, or BMP5-8 (Adomako-Ankomah and Ettensohn, 2013; Duloquin et al., 2007; McIntyre et al., 2013; Piacentino et al., 2016a, 2016b, 2015).

To determine the time dependence of EtOH-mediated defects, first, we treated embryos at different time points ranging from fertilization to 30 hours post-fertilization (hpf), then scored the resulting pluteus larvae for patterning defects at 48 hpf. The results show that an inflection point occurred between 22 and 24 hpf when the fraction of perturbed embryos dramatically declined (Fig. 1D1, left; Fig. S1A, S1B1-4). This indicates that EtOH is most effective before 24 hpf. This timepoint overlaps with both 1° and 2° skeletal patterning processes, in keeping with the broad skeletal defects induced by EtOH (Fig. 1A-C), although 1° skeletogenesis is well-underway at 24 hpf (Piacentino et al., 2016b, 2016a, 2015). Next, to define when EtOH begins to have an effect, we treated embryos at fertilization, removed EtOH hourly over a range of time points, then scored the resulting larvae at pluteus stage. We found an inflection point at 5 hpf, indicating that EtOH effects initiate between 4 and 5 hpf (Fig. 1D2; Fig. S1A, S1B5-6). This time point is surprisingly early, before the onset of Nodal expression and ectodermal dorsal-ventral specification (Bradham et al., 2006). These results indicate that EtOH perturbs skeletal patterning between 5 and 24 hpf.

Because EtOH treatment caused a broad range of skeletal patterning defects, we evaluated whether any defects predominated for different intervals of EtOH exposure during the overall window of sensitivity to EtOH by scoring skeletal defects in embryos treated with EtOH for the same defined temporal windows. Our results show that 2° elements and rotational defects are most sensitive to EtOH at 17 hpf, while 1° defects are most penetrant at 18 hpf (Fig. S1C1). We used the same approach to assess EtOH wash-out embryos, and the results show that 2° elements and rotational defects are most sensitive to EtOH after 6 hpf (Fig. S1C2). These data demonstrate that EtOH is most effective between 7 and 17 hpf regarding specific perturbations to the 2° elements and rotational defects, while impacts on the 1° elements are most pronounced after 18 hpf.

### EtOH indirectly impacts the PMCs to produce skeletal patterning defects

To determine whether the EtOH-mediated defects arise from perturbation of the ectoderm or the PMCs, we performed PMC transplantation experiments (Armstrong et al., 1993; Ettensohn and McClay, 1986; Piacentino et al., 2016b) to produce chimeric embryos in which either the PMCs or the remaining hulls were treated with EtOH (Fig. 1E1). When we transplanted EtOH-treated PMCs into control hulls, this resulted in embryos with normal skeletons (Fig. 1E2). This outcome shows that the skeletal patterning defects do not arise from a direct effect of EtOH on the PMCs, and that the microsurgical manipulations of the hulls did not perturb the skeletal patterning process. In contrast, when we performed the reciprocal experiment in which control PMCs were transplanted into EtOH-treated hulls, the embryos developed skeletal patterning defects characteristic of EtOH treatment (Fig. 1E3). These results demonstrate that EtOH acts on PMCs indirectly via impacts on other tissues such as the ectoderm. This is consistent with EtOH treatment perturbing the expression of skeletal patterning cues rather than directing perturbing the PMCs.

### EtOH does not perturb ectodermal DV specification

The ectoderm is the source of most skeletal patterning cues (Piacentino et al., 2016b). This tissue is subdivided into dorsal and ventral regions during early development by TGF-ß signaling (Duboc et al., 2004); DV perturbations produce radialized embryos and skeletons (Armstrong et al., 1993; Bradham and McCLay, 2006; Hardin et al., 1992; Piacentino et al., 2016a, 2015). While the patterning defects elicited by EtOH do not resemble radialization, we nonetheless tested whether DV specification of the ectoderm is affected by EtOH treatment. We performed immunostains to examine the ectodermal ciliary band (CB), a distinct region that is spatially restricted to the boundary between the dorsal and ventral ectodermal regions by the DV specifying TGF-ß signals in sea urchin embryos. When those signals are disrupted, the ciliary band is either posteriorly positioned or spatially unrestricted (Bradham et al., 2009; Duboc et al., 2004; Yaguchi et al., 2010). Our results show that the CB was both restricted normally and appropriately positioned within the context of the perturbed morphology induced by EtOH treatment (Fig. S2A1. A4). These results suggest that EtOH exposure does not affect ectodermal DV specification. This finding was corroborated by evaluating gene expression for the dorsal marker Lv-IrxA and ventral marker Lv-Chd via single-molecule FISH. The spatial expression of both genes was normal in EtOH-treated embryos (Fig. 2A), confirming that EtOH exposure does not perturb ectodermal DV specification.

**Figure 2.**
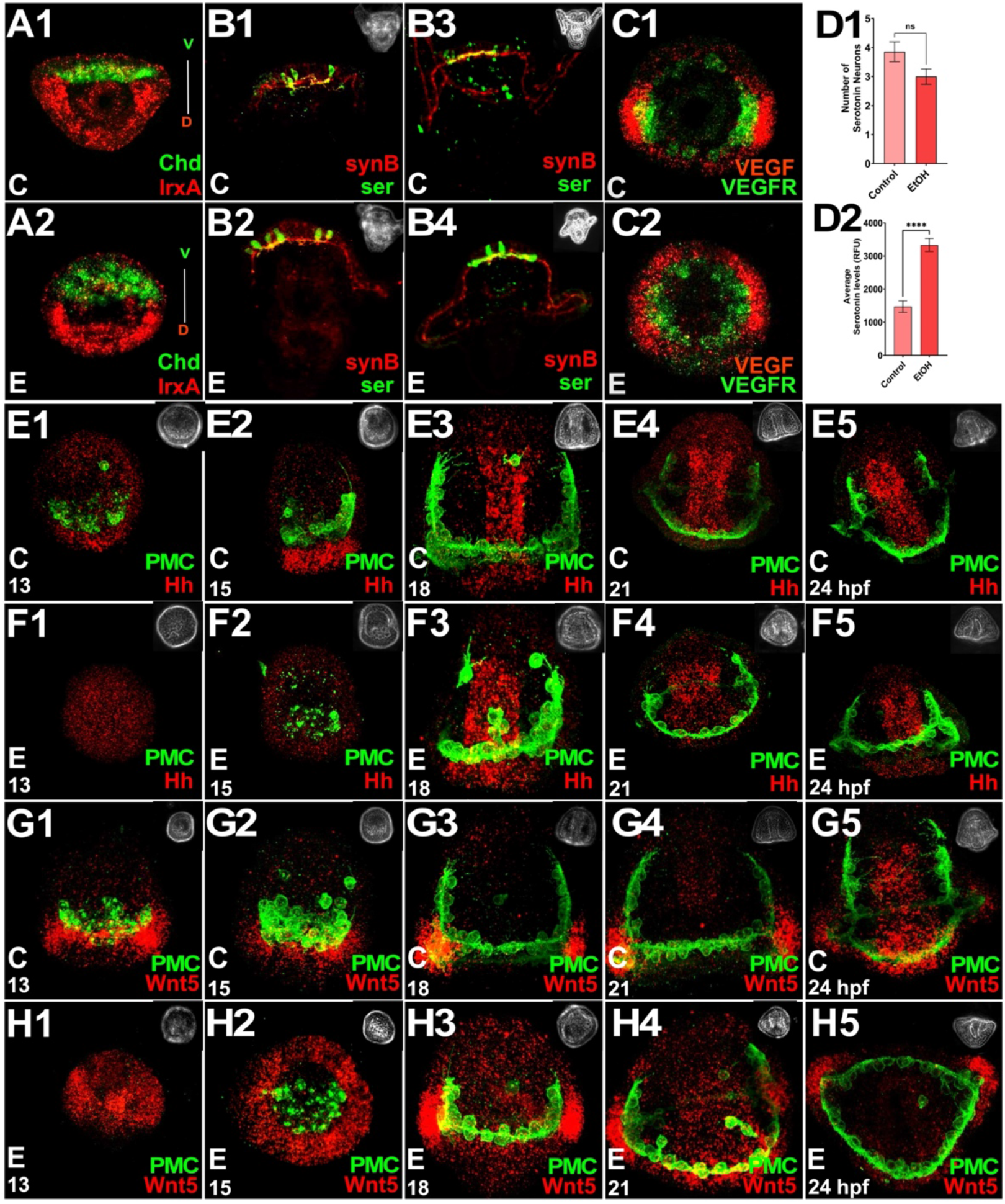
EtOH treatment does not inhibit ectodermal DV specification or neural development and delays normal expression of the PMC-directing signal Wnt5. **A.** The ciliary band was visualized via immunostaining in controls (A1) and EtOH-treated (A2) embryos at 48 hpf. **B.** Neural-specific immunostaining for serotonin (ser, green) and pan-neural synaptotagmin B (synB, red) is shown in control (B1) and EtOH-treated (B2) anterior structures at 48 hpf (pluteus stage, 600x). Embryos are also shown at 100x magnification to capture their complete morphology (B3-B4); corresponding phase contrast images are inset. **C.** HCR-FISH for PMC-specific VEGFR (green) and ectodermal VEGF (red) gene expression is shown in control (C1) and EtOH-treated embryos (C2) at 18 hpf (late gastrula stage); embryos are shown in vegetal views. **D.** The number of neurons (D1) and the level of the serotonin signal per cell (D2) is shown as the average per embryo ± s.e.m.; n = 15 embryos per condition; ****, p ≤ 0.0001 (student t-test). **E-H.** Embryos were collected at the indicated time points then subjected to fluorescence *in situ* hybridization for Lv-Hh (A-B) or Lv-Wnt5 (C-D) along with PMC immunostaining in control (1) and EtOH-treated (2) embryos. Embryos are shown with anterior oriented upward, except E5, F4, F5, and G5, shown with ventral upward. See also Fig. S2-3.

### EtOH does not inhibit neural development

EtOH is a well-known neural teratogen that leads to intellectual impairment and reduced brain size in vertebrates and humans (Hoyme et al., 2005; Khalid et al., 2014). To test whether neural defects are a conserved response to embryonic EtOH exposure, we visualized synaptotagmin B (synB)-positive and serotonergic neurons using immunostaining (Fig. 2B1-B4). Surprisingly, we did not detect any losses of serotonergic neurons in the EtOH-treated embryos (Fig. 2B, 2D1), nor any apparent losses of synB neurons (Fig. 2B). Instead, we found a statistically significant increase in the serotonin immunofluorescence signal in EtOH-treated embryos compared to controls (Fig. 2B, 2D2). Thus, unlike vertebrates, we did not detect an overt loss of neurons with EtOH. Taken together, these results indicate that EtOH exposure does not disrupt neural specification or patterning in sea urchin embryos but strongly increases the serotonin expression level within the nervous system.

### EtOH perturbs spatiotemporal expression of Wnt5 ectodermal patterning cue

The ectoderm is required for normal PMC positioning and biomineralization since it provides instructive cues that are detected by the PMCs (Armstrong et al., 1993; Duloquin et al., 2007; Ettensohn and McClay, 1986; McIntyre et al., 2013; Piacentino et al., 2016a, 2016b, 2015; von Ubisch, 1937). Thus, our next step was to ask whether EtOH exposure perturbs the expression of ectodermal skeletal patterning cues, including Lv-Wnt5 and Lv-VEGF. Lv-VEGF is an ectodermal cue that is expressed in the ectoderm that overlies ventrolateral PMC clusters at the late gastrula stage (Fig. 2C1, red; Fig. S2B1) and is required for normal PMC cluster formation and biomineralization; at later time points, VEGF is required for skeletal patterning (Adomako-Ankomah and Ettensohn, 2013; Duloquin et al., 2007; Piacentino et al., 2016b). Lv-Wnt5 is also expressed in the ectoderm that overlies the PMC clusters (Fig. S2C1) and is required for biomineralization (McIntyre et al., 2013). We found that the expression of both Lv-VEGF and Lv-Wnt5 appears to be spatially expanded to an abnormally broad ectodermal expression pattern in EtOH-treated embryos, with VEGF expression expanded laterally, and Wnt5 expression expanded both laterally and apically (Fig. 2C2 red; Fig. S2B2, S2C2). We also assessed the expression pattern for Lv-Hh, which is expressed by the endoderm and contributes to skeletal patterning (Walton et al., 2009). Since gastrulation is delayed by EtOH treatment, the morphology of the developing archenteron differs from controls such that the gut tube is shorter and wider, making a qualitative assessment of Hh expression domain size challenging (Fig. S2D). Quantitation of their expression areas shows that among these three skeletal patterning cues, only Wnt5 expression is significantly spatially perturbed by EtOH exposure at 18 hpf (Fig. 2J).

To test whether the differences in gene expression might reflect a developmental delay, we compared their expression over time. Those results show that Lv-Hh and Lv-Wnt5 expression patterns match controls with a 3-to 6-hour delay (Fig. 2E-H). Hh expression appears mildly delayed with EtOH treatment compared to controls (Fig. 2E-F). Wnt5 exhibits broad ectodermal expression in controls at 13 and 15 hpf, spatially contracts to bilateral posterior ventrolateral regions of expression at 18 and 21 hpf, then displays a distinct dorsal expression domain at 24 hpf (Fig. 2G). In EtOH-treated embryos, Wnt5 expression similarly contracts to the posterior VL ectoderm at 21 hpf, 3 hours later than controls, while simultaneously exhibiting the dorsal expression domain, 3 hours earlier than controls (Fig. 2G-H). Taken together, these results show that the patterning cues Hh and Wnt5 exhibit temporally abnormal spatial expression in EtOH-treated embryos. While this change to Wnt5 might be responsible for the perturbations to skeletal patterning that occur with EtOH exposure, this seems less likely to account for the patterning defects since Wnt5 loss of function blocks biomineralization rather than patterning per se (McIntyre et al., 2013). The effects on Hh are more likely to contribute to patterning defects since Hh signaling is required for normal 2° skeletal patterning (Walton et al., 2009).

### EtOH delays PMC migration, positioning, and spatial gene expression

Given the effects of EtOH on Wnt5 and Hh expression, we next examined the skeletogenic PMCs in EtOH-treated embryos using immunostaining. PMCs in control embryos ingress at the mesenchyme blastula stage (~13 hpf) (Fig. 3A1), become organized into the 1° ring-and-cords pattern during gastrulation (~15-18 hpf) (Fig. 3A2–A3), then initiate biomineral secretion in the ventrolateral PMC clusters at late gastrula stage (~18 hpf) (Fig. 3A3). In EtOH-treated embryos, PMC ingression and migration were delayed by approximately 3 hours compared to controls, although the cells eventually adopted relatively normal spatial positions by 24 hpf (Fig. 3B1–B5).

**Figure 3.**
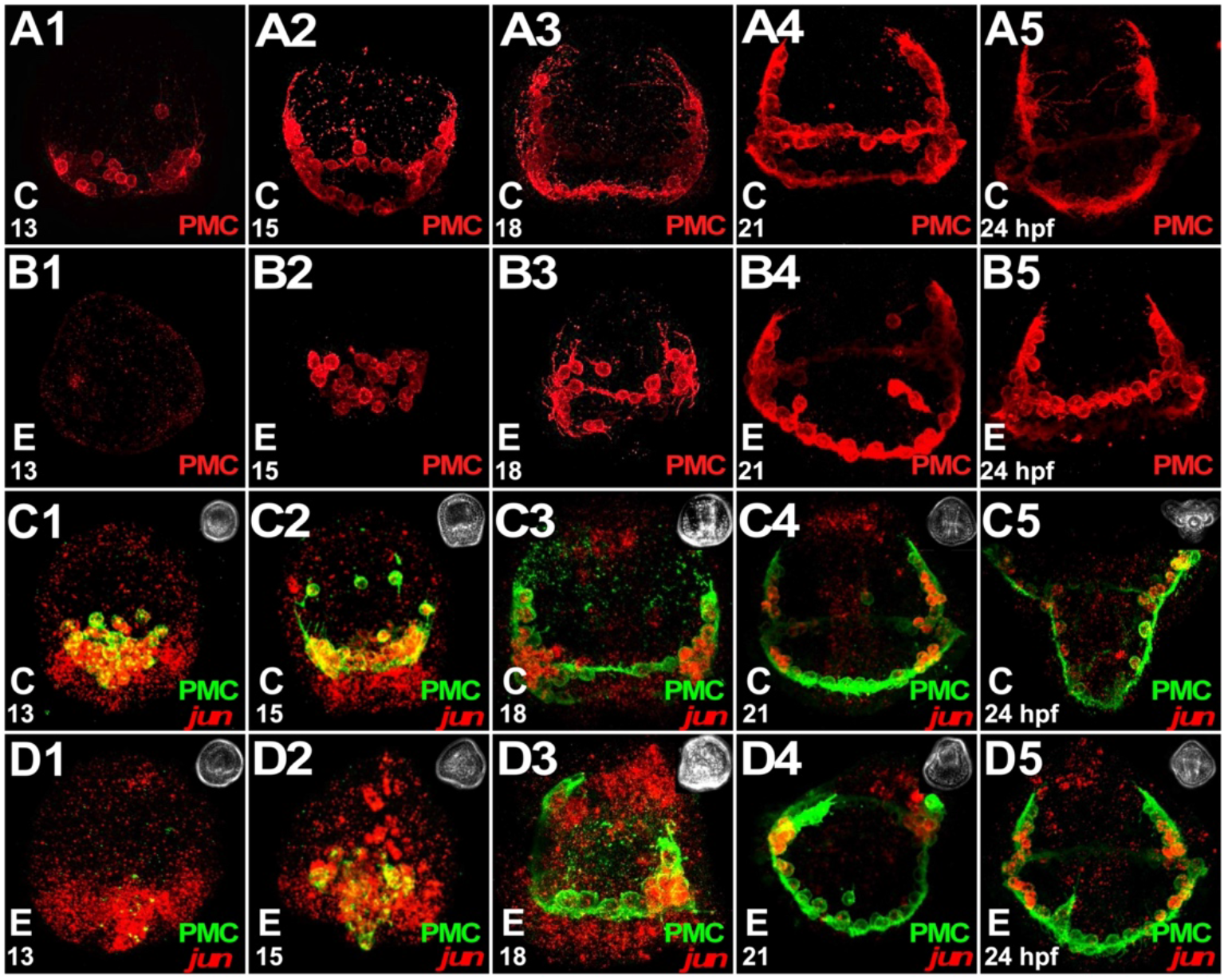
EtOH treatment results in delayed PMC ingression, migration, and Jun expression. **A-B.** PMCs were immunolabeled in control (A) and EtOH-treated embryos (B) that were fixed at 13 (1), 15 (2), 18 (3), 21 (4) or 24 (5) hpf during gastrulation. All embryos are oriented with anterior upward. **C-D.** Lv-Jun expression was detected using FISH and PMCs were immunolabeled in control (1), and EtOH treated (2) embryos at indicated time points. All embryos are oriented with anterior upward except D4 and D5, shown in vegetal views. See also Fig. S2.

During skeletal secretion, PMCs located in distinct regions express different genes (Sun and Ettensohn, 2014; Zuch and Bradham, 2019) reflecting their diversification during skeletal patterning. We next asked whether the spatial expression of known PMC subset genes is perturbed in response to EtOH exposure using a combination of FISH and PMC immunostaining. Lv-VEGFR is generally expressed in the PMCs and is elevated in the PMC clusters at late gastrula stage (Fig. 2C1, green; S2E1), adjacent to the VEGF signal that is expressed in the posterior ventrolateral ectoderm (Fig. 2C1, red) (Adomako-Ankomah and Ettensohn, 2013; Duloquin et al., 2007; Schatzberg et al., 2015; Sun and Ettensohn, 2014). Lv-Jun is a transcription factor that is expressed primarily in the PMC clusters (Fig. 3C3, S2F1-G1) (Sun and Ettensohn, 2014). In EtOH-treated embryos, VEGFR exhibits spatial expanded expression into non-cluster PMCs (Fig. 2C2, green; S2E2), while Jun is not expanded in the PMCs but interestingly is ectopically expressed in the ectoderm at 18 hpf (Fig. SF2-G2). We quantified the expression area for Lv-Jun and found that the spatial domain for Jun is significantly increased by EtOH exposure at 18 hpf (Fig S2J). We also verified the spatial expansion of VEGFR by counting the number of PMCs that expressing VEGFR and found that EtOH-treated embryos exhibit a statistically significant increase in the number of VEGFR-positive PMCs (Fig. S2K). Finally, we also assessed two additional genes enriched in the VL cluster PMCs, Lv-Frp and Lv-Otop. Neither Frp nor Otop exhibited changes in spatial expression with EtOH exposure (Fig.S2D-E, S2K). This observation of perturbed spatial expression of some PMC subset genes is consistent with the perturbation of some patterning cues and agrees with our previous results, suggesting that the EtOH-induced skeletal defects reflect perturbations to patterning cues.

Given that PMC ingression and migration are delayed in response to EtOH, and that patterning cue expression is also delayed, we next evaluated how Jun expression changes with time. The results show that ectodermal expression of Jun normally occurs at 15 hpf in control embryos, while a similar expression pattern was observed in EtOH-treated embryos at 18 hpf, consistent with the general delay in PMC development with EtOH (Fig.3C-D). As with migration, Jun expression appears normally restricted to the PMC clusters in EtOH-treated embryos at later time points (Fig. 3D4-5). Together, these results show delays induced by EtOH exposure in the spatial expression of ectodermal and endodermal patterning cues and a PMC subset gene, as well as delayed PMC ingression, migration, and differentiation.

### EtOH-mediated skeletal patterning defects arise independently of retinoic acid perturbation

A central mechanism that has been identified for FAS phenotypes in vertebrates is the loss of retinoic acid (RA) production and signaling (Kot-Leibovich and Fainsod, 2009; Nelson et al., 2013). This occurs because EtOH is normally metabolized to acetaldehyde by alcohol dehydrogenase (ADH); acetaldehyde then competitively binds and inhibits retinaldehyde dehydrogenase (RALDH), causing the reduced synthesis of retinoic acid (RA) (Shabtai et al., 2018) (Fig. 4A, blue). Experiments performed in zebrafish embryos demonstrated that cotreatment of EtOH and RA was sufficient to rescue the EtOH phenotype (Marrs et al., 2010). In other studies with in frog embryos, EtOH exposure provokes limb defects that are hallmarked by altered RA signaling and rescued by exogenous RA (C. S. Johnson et al., 2007; Yelin et al., 2005). Therefore, to test whether this mechanism is conserved in sea urchins, we first examined whether acetaldehyde is sufficient to phenocopy EtOH. We found that treatment with acetaldehyde caused skeletal patterning defects that are similar but not identical to the defects produced by EtOH (Fig. S3A-C, S3E). Acetaldehyde embryos also showed a similar delay in PMC migration and positioning (Fig. S3C3-C5). We also tested the time dependence of the effects of acetaldehyde and found that the inflection point is delayed compared to EtOH to 25 hpf (data not shown), consistent kinetically with a model in which EtOH perturbs sea urchin development by increasing acetaldehyde levels that result in diminished RA levels, as in vertebrates.

**Figure 4.**
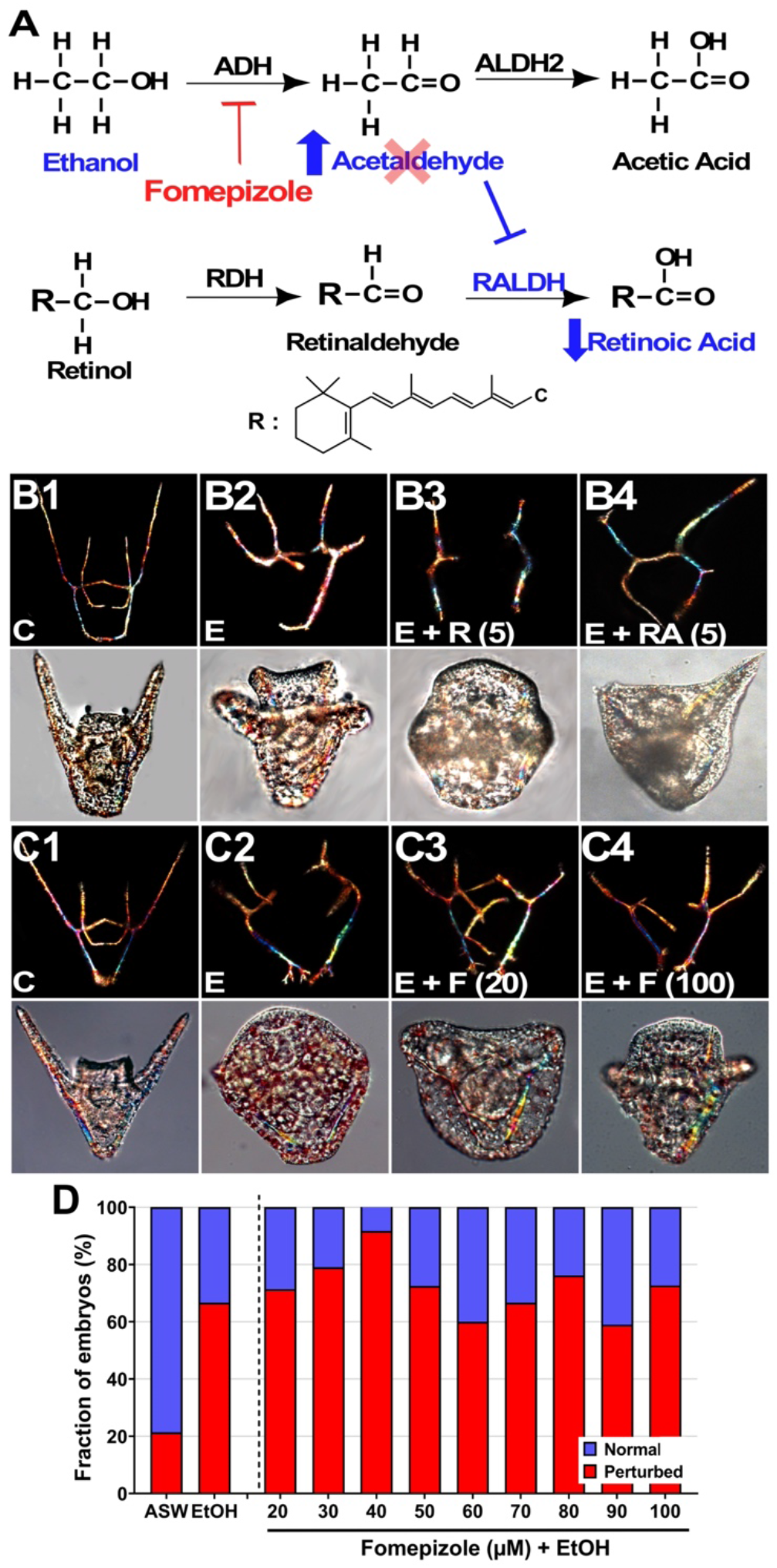
EtOH-mediated defects do not arise via reduction in retinoic acid. **A.** EtOH metabolism results in reduced RA synthesis via acetaldehyde-mediated inhibition of RALDH (blue). Fomepizole inhibits ADH and thereby reduces acetaldehyde production from EtOH (red). **B-C.** Skeletons (bifringence, upper panels) and morphology (DIC, lower panels) are shown at 48 hpf for embryos subjected to treatments with 5 μM Retinol (R) or Retinoic Acid (RA) with or without EtOH (E), or with EtOH (E) and Fomepizole (F) at 20 μM or 100 μM. **D.** The fraction of perturbed embryos is plotted for the indicated treatments; n ≥ 100 embryos per condition in three replicates. ADH, alcohol dehydrogenase; ALDH2, Aldehyde dehydrogenase; ASW, artificial sea water; F, fomepizole; R, retinol; RA, retinoic acid; RDH, retinol dehydrogenase; RALDH, retinaldehyde dehydrogenase. See also Fig. S4.

To test this model, we performed two key experiments. In the first set of experiments, we attempted to rescue the EtOH phenotype by adding RA or retinol at fertilization. We tested a range of doses and found surprisingly that neither RA nor retinol rescued EtOH (Fig. 4B, Fig. S3D). Next, we attempted to rescue EtOH with fomepizole (4-methylpyrazole), an inhibitor of ADH that blocks acetaldehyde production (Fig. 4A, red). We validated the inhibitory activity of fomepizole on EtOH-induced acetaldehyde production in sea urchin embryos (Fig. S3F). However, we found that fomepizole treatment did not rescue the EtOH phenotype over a wide range of doses (Fig. 4C-D), indicating that acetaldehyde production is not required for the defects produced by EtOH. This result is consistent with the lack of rescue of the EtOH phenotype by either RA or retinol. Taken together, these unexpected findings indicate that while acetaldehyde is sufficient to perturb skeletal patterning, neither acetaldehyde production, ADH activation, nor RA reduction account for the mechanism by which EtOH exposure produces skeletal patterning defects in sea urchin embryos. Thus, the role for the RA pathway in vertebrate EtOH teratogenesis is not conserved throughout deuterostomes.

### Inhibition of the Hh signaling pathway partially rescues the EtOH phenotype

The Hh signaling pathway is causally implicated downstream from EtOH in FASD in vertebrate embryos (Abramyan, 2019; Eberhart and Parnell, 2016; Hong and Krauss, 2017; Li et al., 2007; Sidik et al., 2021). Since Hh signaling is also implicated in sea urchin skeletal patterning (Walton et al., 2009), we next asked whether perturbed Hh signaling can explain the EtOH-mediated defects by testing whether the Hh pathway inhibitor cyclopamine or agonist SAG (Chen et al., 2002b, 2002a; Frank-Kamenetsky et al., 2002) can rescue EtOH-treated embryos. Neither cyclopamine nor SAG alone elicited patterning defects (Fig. 5A2-3). When combined with EtOH, cyclopamine provided a modest but statistically significant rescue, while SAG treatment did not. (Fig. 5A4–6, 5B). Interestingly, increasing the dose of cyclopamine did not improve the rescue effect, and cyclopamine was the least effective at the highest tested dose (Fig. S4). Similarly, higher doses of SAG did not improve the rescue effect (Fig. S4). These results show that antagonism but not agonism of Hh signaling is sufficient to partially rescue the EtOH phenotype, but only at low doses and with a weak effect.

**Figure 5.**
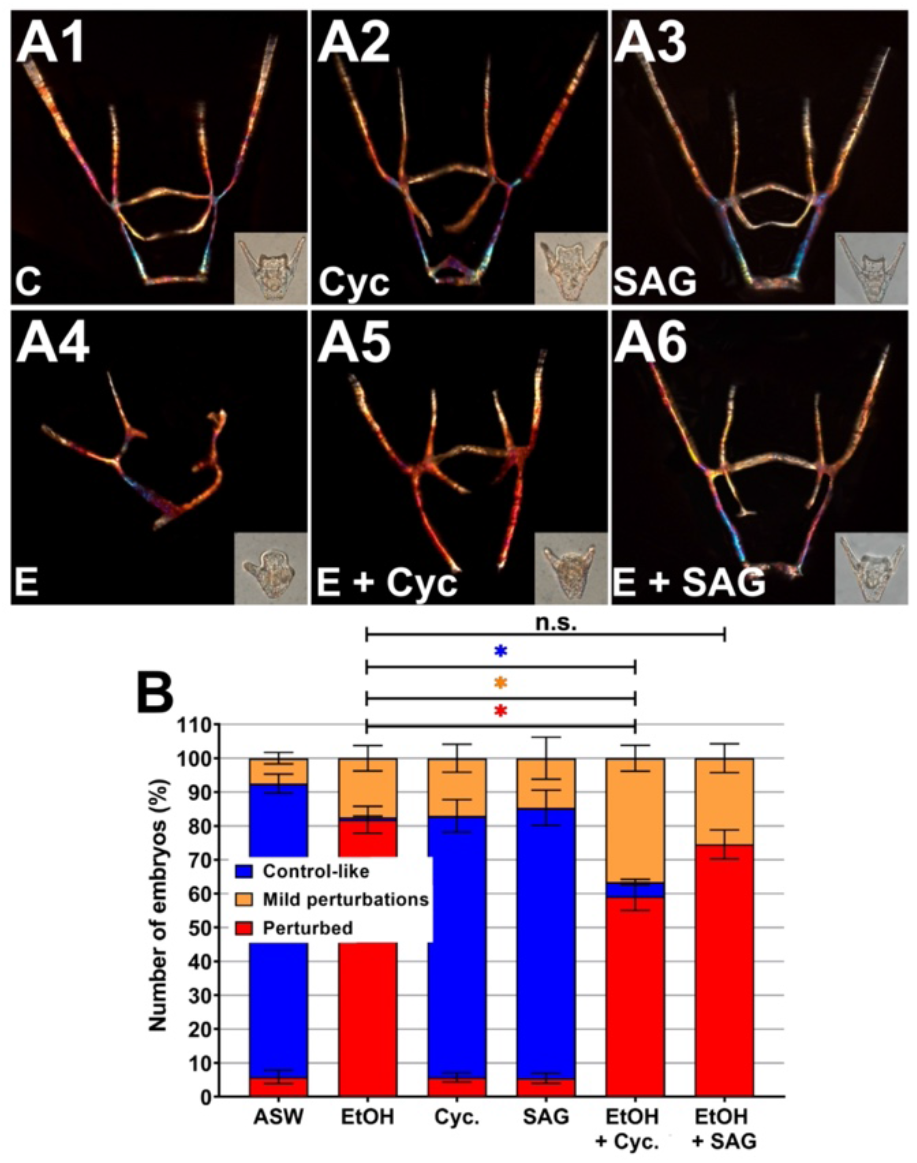
Inhibition of Hh signaling partially rescues the EtOH phenotype. **A.** Skeletal birefringence and morphology (DIC, inset) are shown at 48 hpf for embryos treated with cyclopamine (cyc, 0.2 μM) or with SAG (0.3 μM) with or without EtOH (E). **B.** The fraction of normal and perturbed embryo is plotted for the indicated conditions as the average ± s.e.m.; n ≥ 150 embryos per condition in three replicates; * p < 0.05; n.s., not significant (t test). Mild perturbations are illustrated by panels A5 and A6. See also Fig. S5.

**Figure 6.**
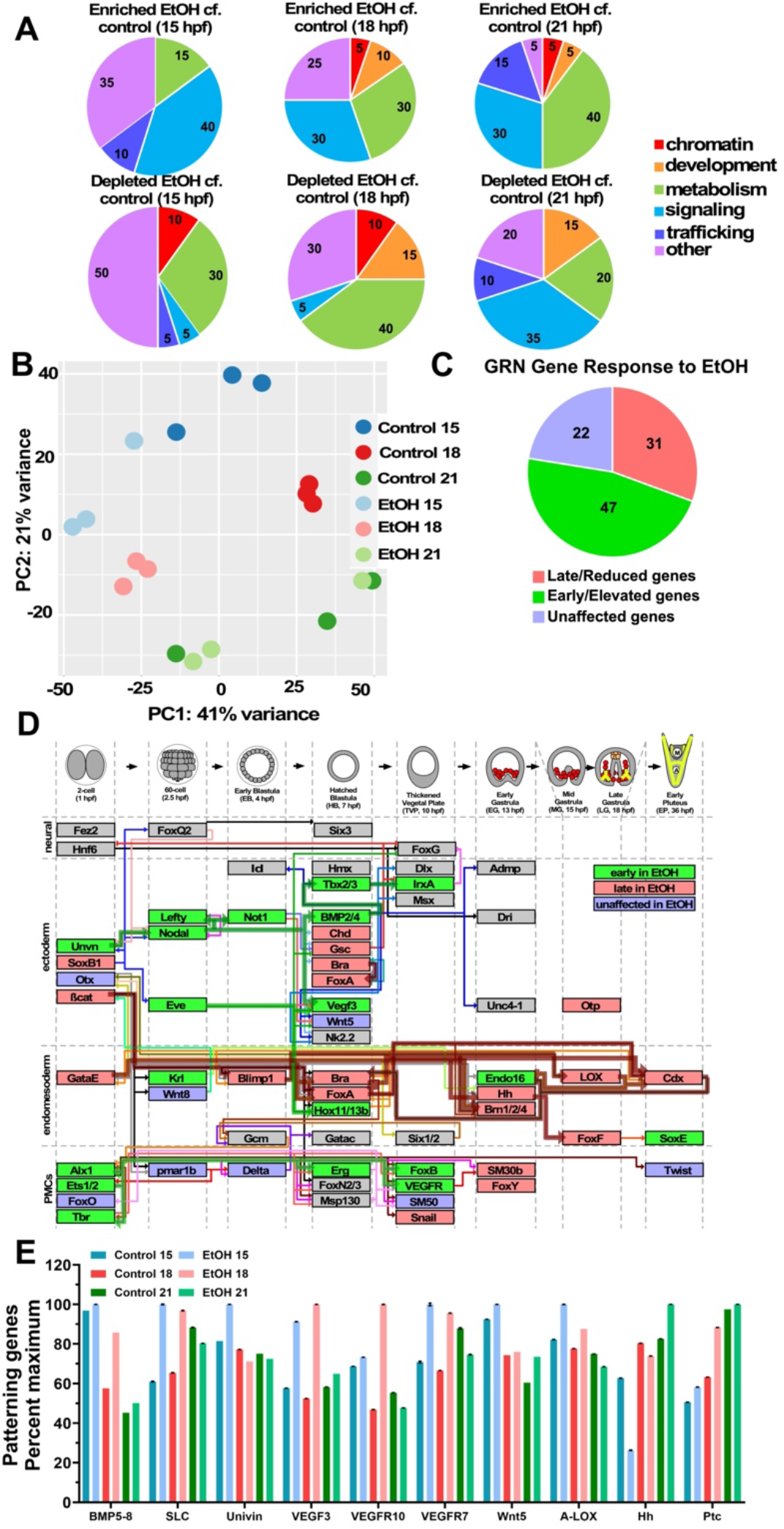
EtOH treatment results in disrupted temporal synchrony. **A.** Broad categories for the top 20 GO terms enriched or depleted in EtOH-treated embryos compared to time-matched controls are shown; their relative percentages are indicated. **B.**2-D principal component analysis (PCA) compares control and EtOH-treated replicates. **C.** The effects of EtOH on the expression of 49 GRN genes at three time points is shown by their distribution into late/reduced, early/elevated, and unaffected bins; percentages are indicated. **D.** An overview of the sea urchin specification GRN model is separated vertically into major tissues, and horizontally by time, with genes positioned according to their onset of expression (adapted from Hogan 2020). Colored nodes indicate late or reduced genes (red), early or elevated genes (green), and unaffected genes (blue). Plausible circuit connections are indicated as thick, transparent edges. **E.** The expression levels of genes encoding known patterning cues in control and EtOH-treated embryos at 15, 18, 21 hpf are shown as the average ± s.e.m. with values scaled to the maximum expression level for each gene. See also Fig. S5-6.

### RNA-seq reveals temporal perturbations of GRN gene expression

We used RNA-seq to query the global effects of EtOH on development by comparing the transcriptomes of control and EtOH-treated embryos at 15, 18, and 21 hpf. We found many differentially expressed (DE) genes at 15 and 18 hpf, but at 21 hpf, far fewer DE genes were identified (Fig. S5A). We analyzed differential GO pathway activity using ssGSEA to identify functional categories of genes affected by EtOH exposure. The top 20 enriched and depleted GO terms for each pair-wise comparison were binned into broad categories (Fig. 6A, S5B). Those results show that metabolism and signaling were the GO term categories most impacted in EtOH-treated embryos in time-matched comparisons, accounting for 55-70% of the enriched and 35-55% of depleted GO terms (Fig. 6A). Heterochronic comparisons revealed additional differences in chromatin and protein trafficking while still exhibiting a majority of differences in signaling and metabolism (Fig. S5B).

The first two components of a principal component analysis (PCA) accounted for 62% of the overall variation in the data. PC1 appears to sort on treatment for the earlier time points but does not separate control and EtOH at 21 hpf (Fig 6B), consistent with the few DE genes identified at 21 hpf. PC2 appears to sort on time, with EtOH samples being temporally offset from controls, aside from 21 hpf. Given the mild temporal displacements in the PCA results and the conspicuous morphological delays in development for EtOH-treated embryos, we further assessed EtOH-mediated temporal perturbations by comparing the expression of genes in the known gene regulatory network (GRN) specification models for sea urchins, focusing on genes whose expression changes between 15 and 21 hpf in *L. variegatus* embryos and their regulators (Davidson et al., 2002; Hogan et al., 2020; Li et al., 2014; Materna et al., 2013; Peter and Davidson, 2011; Saudemont et al., 2010; Su et al., 2009), using a cut-off threshold of ≥ 18% change. Surprisingly, of the 49 genes we evaluated, only 30% exhibited late or reduced expression, with 47% of genes instead displaying early or elevated expression, while the remaining 22% were unaffected by EtOH treatment (Fig. 6C). Temporally perturbed genes were present in each of the germ layers (Fig. S6).

These results suggested that some GRN subcircuits are independently activated and therefore insulated from other circuits. To obtain a clearer view, we mapped the late/reduced genes and the early/elevated genes onto a network model and indicated previously known connections that could potentially integrate these changes. Several plausible circuits emerged from this analysis, including elevated or early expression of ectodermal Univin, Nodal, Lefty, Not1, Vegf3, BMP2/4, Tbx2/3, and IrxA (Fig. 6D). Curiously, Nodal is known to activate Lefty, Chordin (Chd), Goosecoid (Gsc), and BMP2/4, but both Chd and Gsc exhibit late or reduced expression rather than early or elevated expression in response to EtOH, implying additional regulators for these genes (Fig. 6D). Similarly, early, or elevated expression interconnects the PMC genes Ets1/2, Alx1, Tbr, Erg, FoxB, and VEGFR. As with the ectoderm, some known targets of Alx1 and Ets1/2 are not affected or are delayed/reduced, including SM50, SM30b, Snail, and Twist (Fig. 6D), once again implying additional unknown regulators for those genes. Interestingly, the clearest circuits interconnecting genes that are delayed or reduced by EtOH are in the endomesoderm and include ß-catenin, GataE, Blimp1, Bra, FoxA, Hh, LOX, and Cdx. The delayed or reduced expression of these genes could explain the delay in gastrulation in EtOH-exposed embryos; perhaps the timing of gastrulation sets the pace for development overall, leading to the morphological delay despite the presence of genes whose expression is early or elevated rather than delayed or reduced. Finally, we evaluated the expression of known patterning genes from the RNA-seq data (Fig. 6E). Those results indicated early or elevated expression for BMP5-8, Univin, SLC, VEGF, VEGFR, and Ptc, late expression for Hh, and unaffected expression for Wnt5. These findings agree with our FISH results for VEGF and Wnt5, particularly that normal expression is achieved at late timepoints, and suggest that Wnt5 is not significantly perturbed by EtOH. These results also support the weak rescue effects we obtained when Hh signaling was perturbed along with EtOH treatment.

## Discussion

In this study, we show that EtOH exposure is teratogenic to sea urchin embryos; more specifically, EtOH perturbs skeletal patterning but does not inhibit neural development. We further show that the defective skeletal patterns in EtOH-treated embryos arise from perturbation to patterning cues rather than from the direct effects of EtOH on the PMCs during skeletogenesis, and are not explained by perturbation to the RA or Hh pathways that are implicated in vertebrate fetal alcohol syndrome disorders. Finally, we observed temporal perturbations in developmental GRN gene expression in EtOH-treated embryos that potentially account for their aberrant pattern formation.

Although skeletal patterning defects were observed that could be considered to be analogous to the facial perturbations in FASD, neural specification and development was not blocked in sea urchins after the exposure to EtOH. Studies in mice, fish, and frogs have shown that EtOH exposure disrupts every step of CNS development, including neural proliferation, migration, and differentiation, as well as directly leading to neural apoptotic and necrotic cell death (Blader and Strähle, 1998; Gil-Mohapel et al., 2019; Sulik, 2014). The absence of evident neural specification or patterning defects in EtOH-treated sea urchin embryos suggests that the neural developmental vulnerability to ethanol is a vertebrate novelty whereas the impact of ethanol on skeletal patterning is more deeply conserved among deuterostomes.

Although the number of serotonergic neurons was not affected by EtOH treatment, the levels of serotonin expression per cell were strongly and significantly increased. This is a surprising outcome. However, because the function of those neurons remains unknown, we are unsure how to interpret the serotonin increase. It will thus be of interest to test the effects of serotonin perturbation in future studies.

Previous studies have linked FASD abnormalities, both neural and facial, to perturbations of the Hh and RA signaling pathways (Ahlgren et al., 2002; Cayuso et al., 2006; Cohen and Sulik, 1992; Petrelli et al., 2019; Reimers et al., 2004; Smith et al., 2014), each of which plays important roles in the development of both the CNS (Chatzi et al., 2013; Janesick et al., 2015; Li et al., 2021) and the facial skeleton (Abramyan, 2019; Dubey et al., 2018; Gur et al., 2022; Sun et al., 2020). Experiments in zebrafish (Marrs et al., 2010) and frogs (C. S. Johnson et al., 2007; Yelin et al., 2005) demonstrated that exogenous RA suffices to rescue EtOH-mediated neural and skeletal defects; similarly, addition of the Hh pathway agonist rescues development of EtOH-exposed zebrafish (Sidik et al., 2021). While the precise mechanism of these effects remains unclear, embryonic exposure to EtOH can induce holoprosencephaly, strongly implicating an inhibitory effect on Shh signaling (Hong and Krause, 2017); in keeping with this, an ethanol-mediated blockage to Shh protein processing has been reported (Li et al., 2007).

Our results show that RA signaling does not play a role in EtOH-mediated perturbations, and Hh plays only a minor role in sea urchin embryos. We found that acetaldehyde also results in skeletal patterning defects, implying that other downstream pathways that are shared by ethanol and acetaldehyde, but that do not impinge upon RA levels, are responsible for their overlapping patterning phenotypes in sea urchins. These shared downstream effects potentially include metabolic pathways, which clearly are affected by EtOH treatment (Fig. 6A); for example, both EtOH and acetaldehyde exert inhibitory effects on mitochondrial β-oxidation (Latipää et al., 1986; Ontko, 1973; You and Arteel, 2019). Thus, the contributions of the RA and Hh pathways to the phenotypic effects of embryonic EtOH exposure surprisingly are not evolutionarily conserved in sea urchins.

We found that EtOH becomes effective quite early during development, beginning around 5 hpf, which is prior to the skeletal patterning events that occur following 13 hpf. We also find that some PMC specification genes in the GRN are affected by EtOH exposure in that they exhibit perturbed timing and/or expression level. Because PMC specification is autonomous, it is likely that EtOH directly perturbs the expression level or timing of those genes. Similar perturbations throughout the GRN likely explain the early effects of EtOH. However, our experiments with chimeric embryos show that EtOH perturbs skeletal patterning by directly affecting ectodermal patterning cues rather than directly impacting the PMCs. This seeming discrepancy arises because two different events are being considered: early PMC specification, and later skeletal patterning. These two processes are experimentally separable since chimeric embryos are produced using PMCs only after their specification is complete, and therefore only query the later patterning process. Thus, our results indicate that EtOH likely directly perturbs PMC specification via changes to the timing and/or levels of expression of PMC specification genes, as well as indirectly perturbing the PMCs during skeletal patterning via direct changes to the timing and/or levels of expression of ectodermal patterning signals that the PMCs receive.

One of the most conspicuous defects in EtOH-treated embryos is a three-hour morphological delay in development. This delay is echoed by the expression of numerous genes, including delays or reductions in the level of expression or the normal spatial pattern for some genes that encode skeletal patterning or differentiation cues, such as Wnt5 and Hh, or PMC subset genes, such as Jun. However, other genes encoding patterning cues exhibit early expression, including BMP5-8, SLC26a2/7, and VEGF3, while still others are unaffected. Temporal discrepancies are present throughout the GRN; those genes exhibit mostly early or elevated expression in response to EtOH, with a minority being unaffected. Together, this suggests that the EtOH-induced phenotype might arise from temporal mismatches between the instructive ectoderm and endoderm tissues and the responsive PMCs, reflecting an overall loss of synchronization across the embryo. Asynchronous intraembryonic development is clearly implied by plausible circuits connecting genes whose expression is early or elevated by EtOH exposure in the ectoderm and PMCs, alongside plausible circuits for genes whose expression is late or reduced by EtOH in the endomesoderm; in each of these tissues, genes with all three temporal profiles are present but not readily explained, implying the existence of still-unknown network components. Temporal loss of registration could also explain why our attempts to rescue the EtOH phenotype have met with limited success since that model suggests a more complex cause for the EtOH-mediated defects than the simple loss of a signal or set of signals. It will be interesting to determine whether these effects are conserved by testing whether EtOH exposure mediates similar temporal disruption in other species.

Overall, our study offers a novel model to study EtOH-mediated teratogenesis that separates the impact on skeletal patterning from the effects on neuronal survival, providing insight into the evolution of the embryonic response to EtOH and revealing a novel dysregulation of temporal synchronization of GRN gene expression that is induced during EtOH-mediated teratogenesis.

## Acknowledgements

We thank Professors Charles Ettensohn, Robert Burke, and David McClay for their gifts of antibodies, and Dr. Todd Blute for microscopy advice. We also thank Prof. Malika Jeffries-El for the use of an oxygen-free glove box and Mr. Leon Novak for his technical assistance. This work was supported by NSF IOS award 1656752 (CAB).

## Author Contributions

This study was conceived by CAB and designed by CAB and NRS. The experiments were executed by NRS, NS, AL, MP, KD, and AEC; bioinformatics analyses were performed by DYH, NRS, and CAB. The manuscript was written by NRS, DYH and CAB, and edited by all co-authors.

