## Supplemental Figures for "Ethanol Exposure Perturbs Sea Urchin Development and Disrupts Developmental Timing"

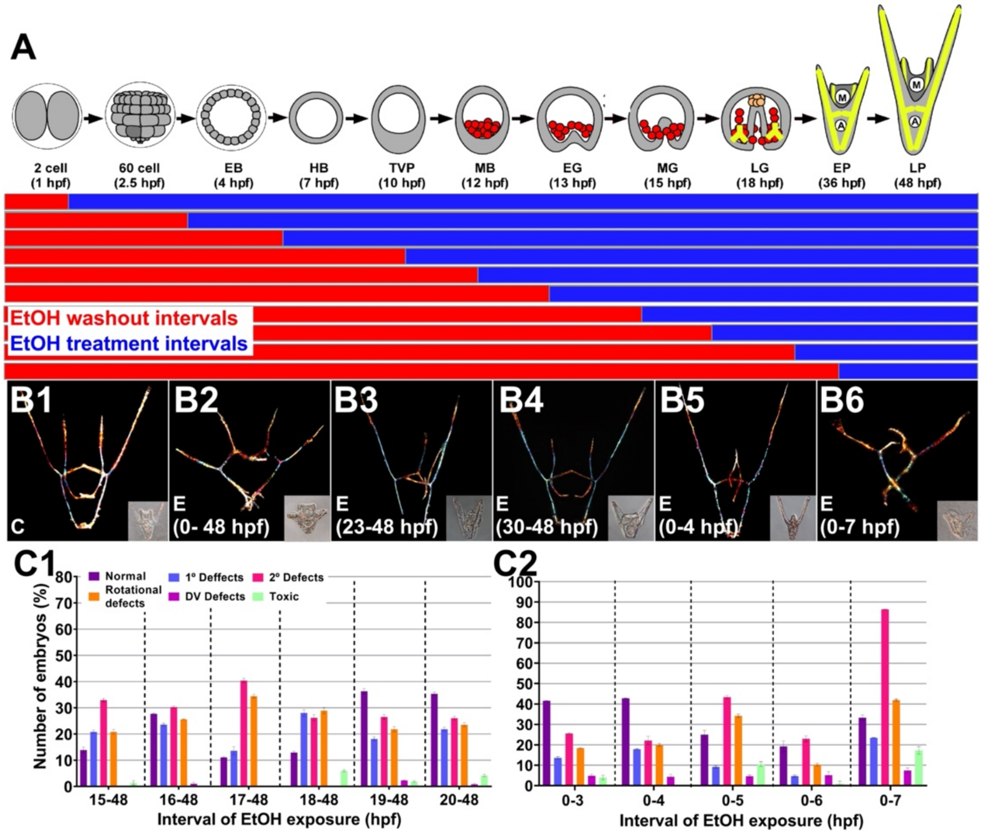


**Figure S1. EtOH is most effective between 5-22 hpf. A.** A schematic of the developmental time course for sea urchin embryos (top) and the temporal treatment regimen (bottom). **B.** Skeletal birefringence and DIC morphology (inset) are shown at 48 hpf for controls (C) and embryos treated with EtOH (E) during the indicated intervals. **C.** Skeletal patterning defects were scored for the indicated groups of EtOH-treated embryos, comparing defects with variable times of EtOH addition (C1) or removal (C2); n ≥ 30 per condition. See also Fig. 1. Panel A was adapted from Hogan et al. 2020.

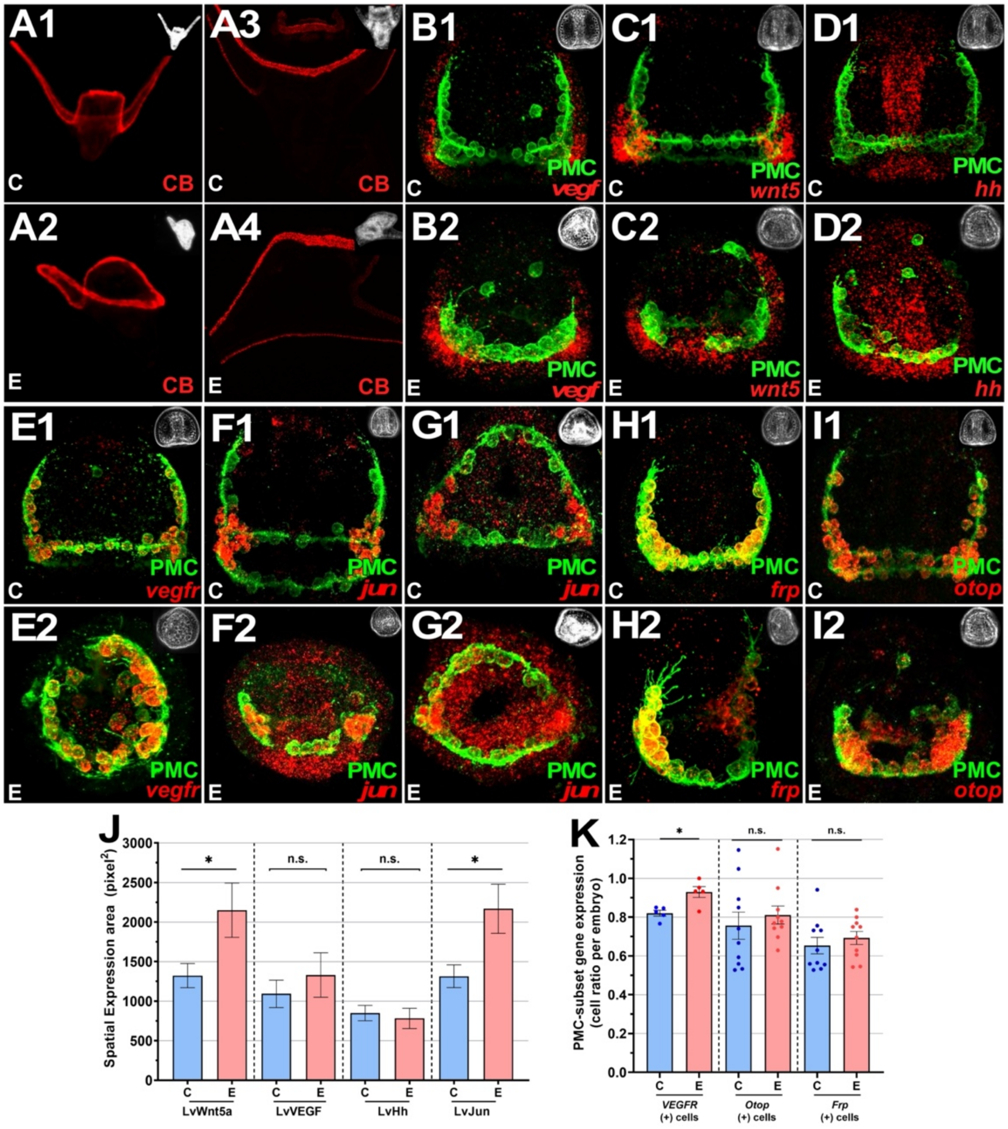


**Figure S2. EtOH treatment results in spatially expanded gene expression of PMC subset genes Jun and VEGFR and ectodermal gene Wnt5 at late gastrula stage but does not perturb ectodermal DV specification. (A-E)** Gene expression was detected using HCR-FISH, and PMCs were immunolabeled in control (1) and EtOH-treated (2) embryos at 18 hpf (late gastrula stage). PMC-specific genes are Lv-VEGFR (A), Lv-Jun (B-C), Lv-Frp (D), and Lv-Otop (E). All embryos are oriented with anterior upward except A2 and C1-2, shown in vegetal views. **F.** The ciliary band (CB) at the DV boundary was immunolabeled in controls (1), and EtOH-treated (2) embryos at pluteus stage (48 hpf) and are shown at 100x (F1-F2) and 600x magnification (F3-F4) to capture their complete morphology. Corresponding phase-contrast images are inset. **G-I.** The expression of the ectodermal signals VEGF (G), Wnt5 (H), and Hh (I) is shown along with PMC immunostaining in control (1) and EtOH-treated (2) embryos at 18 hpf. Embryos are oriented with anterior at the top. **J.** The total expression area for the indicated signals in control (C) and EtOH (E) is shown as the mean summed area per embryo ± s.e.m; *p < 0.05 (student t-test, n=15 per condition). **K.** The proportion of individual PMCs expressing indicated PMC-subset genes were quantified in control and EtOH-treated embryos. Individual counts were plotted as the mean fraction of PMCs that express each gene per embryo ± s.e.m.; n = 10 for Otop and Frp samples and n=7 for VEGFR samples. See also Fig. 2-3.

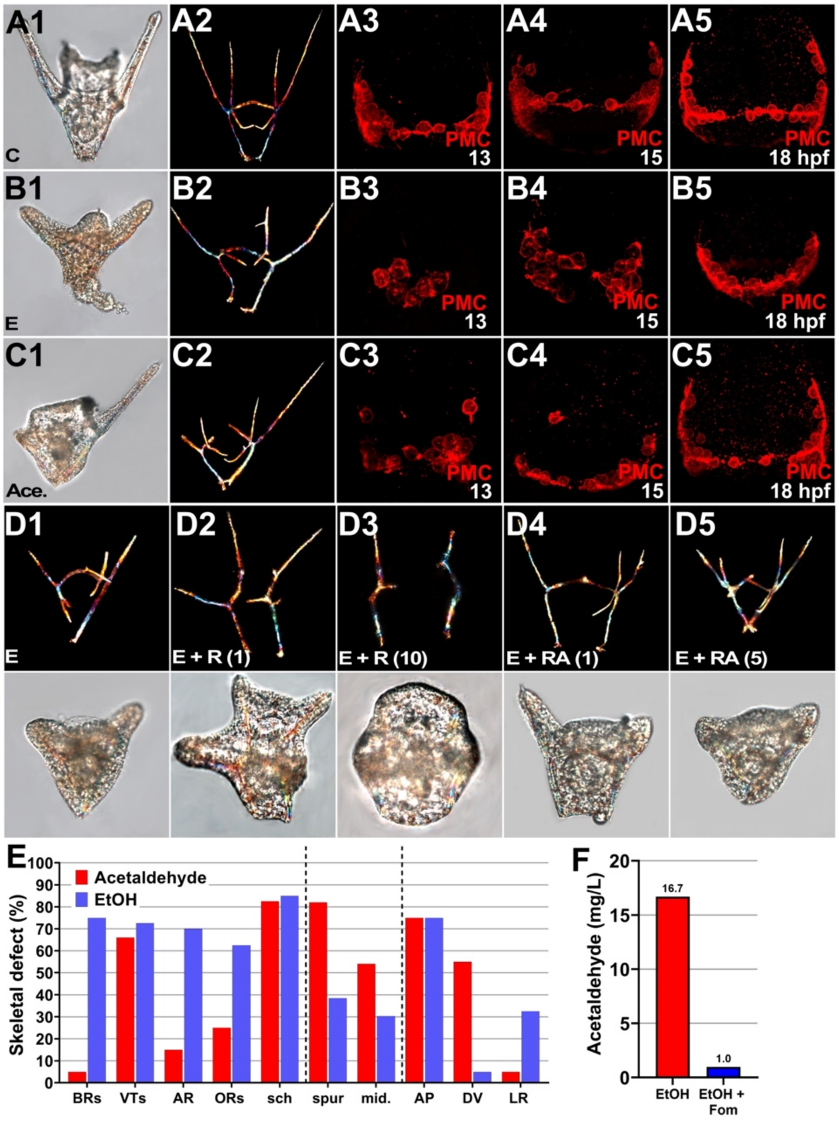


**Figure S3. While acetaldehyde (Ace.) treatment results in skeletal patterning defects, neither retinol (R) or retinoic acid (RA) is sufficient to rescue EtOH. A.-C.** Representative morphology (DIC, 1), skeletal birefringence (2), and PMC immunolabeling are shown at 48 hpf (A, B) or at the indicated time points (C) for control (C, A), EtOH- (E, B, 369 mM), and acetaldehyde- (Ace, C, 891 µM) treated embryos. **D.** Skeletal birefringence (upper panels) and corresponding morphology (DIC, lower panels) is shown at 48 hpf for embryos treated with EtOH (E) and either retinol (R) at 1 µM (D2) or 10 µM (D3), or retinoic acid (RA) at 1 µM (D4) or 5 µM (D5) **E.** Skeletal patterning defects were scored for embryos treated with EtOH or acetaldehyde, including element losses (see Fig. 1A for abbreviations), spurious elements (spur), midline defects (mid) and rotational defects (AP, DV, and LR; see Fig. 1A-B), and are plotted as the percentage of embryos exhibiting the indicated effect; n > 60 embryos per condition**. F.** Acetaldehyde assays confirm the effectiveness of the ADH inhibitor fomepizole (Fom, 100 µM) in sea urchin embryos. The density of embryos for each sample was 1000 embryos/ml. See also Fig. 4.

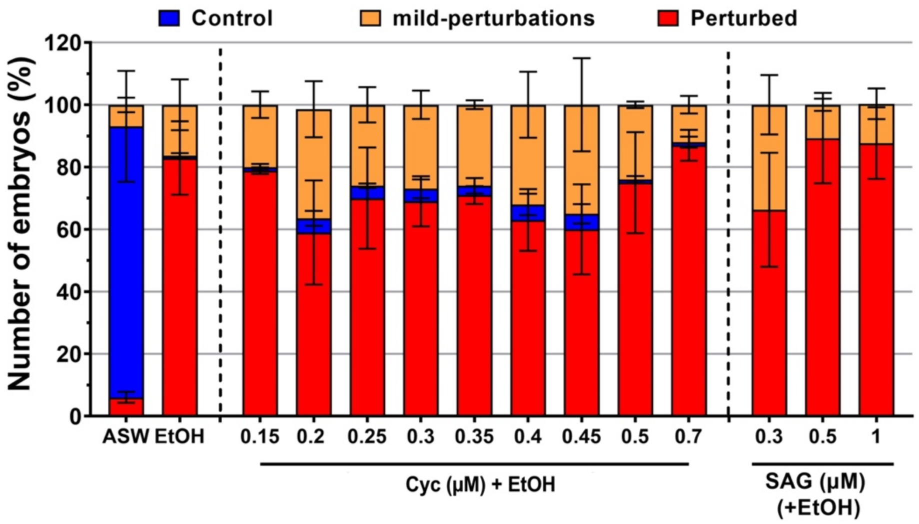


**Figure S4. The rescue of the EtOH phenotype by Hh pathway inhibition is most effective at low doses.** The fraction of skeletal patterning defects are shown for the combinations of EtOH and cyclopamine or SAG at the indicated doses as the average ± s.e.m; n = 150 embryos per condition. See also Figure 5.

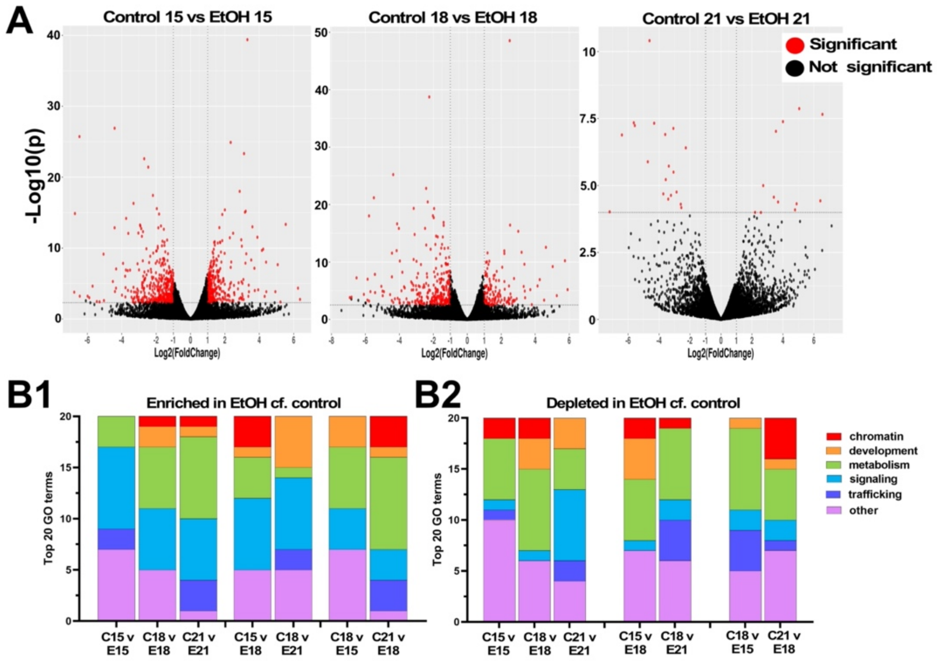


**Figure S5. GO term analysis show metabolism and signaling are the most affected in EtOH samples. A.** Volcano plots show significant DE genes (red) between EtOH and control transcriptomes at each sequenced time point. **B.** The top 20 enriched (B1) or depleted (B2) GO terms, binned into broad categories, are shown for the indicated time-matched or heterochronic comparisons of EtOH to controls. See also Figure 6.

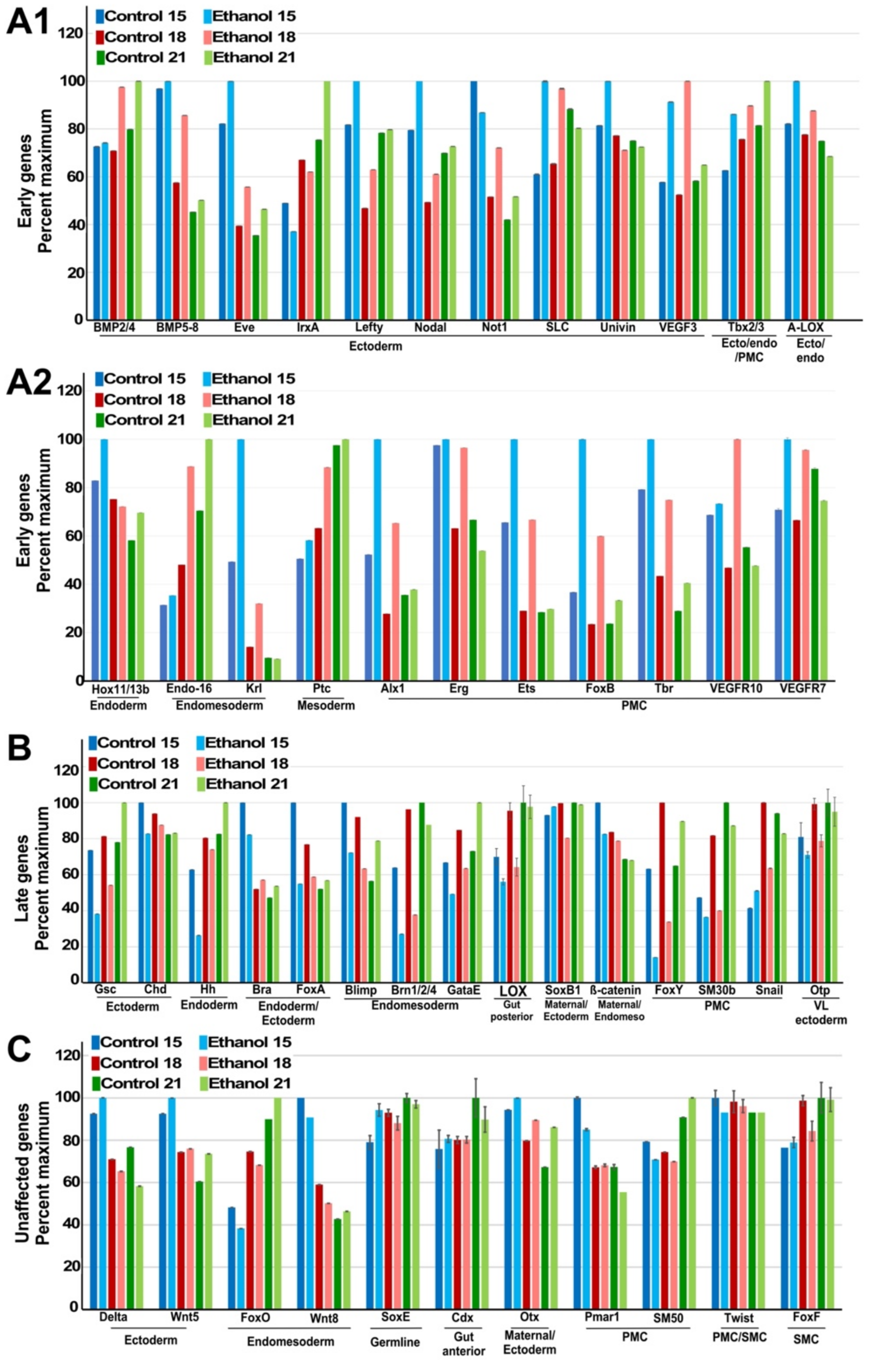


**Figure S6. EtOH-treated embryos exhibit temporally disrupted gene expression.** GRN genes whose expression is either early or elevated (A), late or reduced (B), or unaffected (C) by EtOH treatment are shown as the average ± s.e.m. with expression scaled to the maximum expression level for each gene. The early/elevated group is divided into ectodermal (A1) and endomesodermal (A2) genes. The germ layer in which each gene is expressed is indicated. See also Figure 6.
